# Mechanistic insights into the inhibition of dengue virus NS5 methyltransferase by herbacetin

**DOI:** 10.1101/2024.10.14.618121

**Authors:** Mandar Bhutkar, Shalja Verma, Vishakha Singh, Pravindra Kumar, Shailly Tomar

## Abstract

Herbacetin (HC) is a naturally occurring flavonoid compound with a dual antiviral mechanism. It inhibits the polyamine biosynthetic pathway and targets the methyltransferase (MTase) enzyme of the dengue and chikungunya virus. However, understanding the detailed inhibition mechanism of DENV MTase by HC remains unclear. The crystal structure of DENV3 nonstructural protein 5 (NS5) MTase in complex with HC and S-adenosyl-L-homocysteine presented in this study gives structural insights into the inhibition mechanism. Structural analysis revealed that HC binds to the cap 0 RNA site near the GTP binding site in the DENV3 NS5 MTase, and fluorescence polarization assay demonstrated HC inhibits GTP binding with an inhibition constant K_i_ value of ∼ 0.43 µM. This is the first study that identified an inhibitor that targets the conserved RNA-binding region of NS5 MTase, suggesting its potential as a highly effective scaffold for broad-spectrum antiviral agents against orthoflaviviruses.

**Importance:** Dengue virus (DENV) is a vector-borne disease affecting global health, and no antiviral treatment is available for Dengue fever. DENV NS5 methyltransferase (DENV NS5 MTase) is a viral RNA capping enzyme essential for the virus replication. This study demonstrated that herbacetin (HC), a natural flavonoid, blocked the binding of GTP to NS5 MTase and inhibited the enzyme. In addition to DENV3, HC effectively inhibited the replication of DENV1,2, and 4 serotypes. The crystal structure of the DENV3 NS5 MTase in complex with HC revealed that HC binds at the cap 0 RNA binding site, which is close to the GTP binding site and thereby perturbs the spatial arrangement of GTP binding site residues. HC, as the first identified inhibitor that targets the conserved RNA-binding site of DENV3 NS5 MTase, presents a potential scaffold for the development of new antiviral drugs.

## Introduction

The *Flaviviridae* family comprises the most widespread arboviruses that can cause life-threatening human diseases. This family includes the genus *Orthoflavivirus,* which encompasses various viruses such as Dengue virus (DENV), Zika virus (ZIKV), West Nile virus (WNV), and Japanese encephalitis virus (JEV), etc (1). DENV has four (DENV 1-4) serotypes that can cause infection ranging from asymptomatic to mild symptoms such as fever, rash, joint pain, headache, etc. In severe cases, DENV infection can lead to dengue shock syndrome and dengue hemorrhagic fever (2). DENV gets transmitted by *Aedes aegypti* and *Aedes albopictus* mosquitoes. Approximately 390 million infections occur annually, with most cases reported from tropical and subtropical countries (3). DENV infections are endemic in more than 30 countries, including India, Thailand, Sri Lanka and Myanmar (4). Although the FDA has approved the vaccine for DENV (Dengvaxia), its use is limited to individuals living in endemic areas aged 6 to 16 years with previously laboratory-confirmed DENV infection (5). Considering the limitations of the vaccine and the absence of antiviral drugs, there is an imminent need for the development of effective antiviral treatments for DENV infection.

DENV is an enveloped, positive sense RNA virus having a single open reading frame, which encodes for three structural proteins (capsid, envelope, and preM) and five nonstructural proteins from NS1 – NS5 (6). NS5 is the largest and most highly conserved protein, comprising approximately 900 amino acid residues among all nonstructural proteins (7). It consists of two domains, N-terminal methyltransferase (MTase) and C-terminal RNA-dependent RNA polymerase (RdRp) where each domain can function independently of the other (8). The viral RNA has a type 1 cap structure (m7GpppAmp) at the 5’ end, which plays a crucial role in the translation of viral proteins as well as in evading host immune responses (9). The mechanism of orthoflaviviral MTase comprises two steps. The first step involves the transfer of a methyl group from s adenosine methionine (SAM) to the N7 position of the guanine base situated at the 5’ terminus of the viral RNA. The second step involves the methylation of 2 O’ ribose sugar of the initially transcribed nucleotide (9,10). In orthoflaviviruses, K61-D146-K180-E216 amino acid arrangement is essential for the 2 O’ methylation and remains conserved. It is a catalytic tetrad where Lys180 plays a crucial role in transferring methyl group from SAM to 2 O’ ribose sugar of adenine. Additionally, Lys61, Asp146, and Glu216 help to stabilize the substrate and the transition state during the reaction, ensuring that the 2 O’ methylation occurs efficiently (11). DENV NS5 MTase x-ray crystal structure revealed the presence of s adenosine methionine (SAM), guanosine triphosphate (GTP), and cap 0 RNA binding sites (8,11). Since NS5 MTase has only one SAM binding site and dual methylation function (N7 and 2 O’), the RNA substrate must be repositioned to methylate 2 O’ ribose sugar of the initially transcribed nucleotide (12).

Herbacetin (HC) is a naturally occurring flavonoid in plants, principally flaxseed, parsley, and equisetum (Ramose scouring rush) (13). Herbacetin is structurally close to quercetagatein and isoflavone and exerts various pharmacological activities, including antiviral, antioxidant, anti-inflammatory, and anticancer effects (13–15). Moreover, HC is a known allosteric ornithine decarboxylase inhibitor that depletes cellular polyamine levels upon treatment (16,17). Furthermore, HC exhibited viral MTase inhibition, leading to antiviral activity against DENV and CHIKV with IC_50_ values in µM range (17). However, the structural aspects of inhibition of DENV3 MTase by HC are yet to be revealed and are important to develop a high-efficacy scaffold for the development of drugs. Also, the effect of HC interaction on the binding of GTP in DENV3 NS5 MTase is unknown.

To address these questions, this study presents a detailed crystal structure of the DENV NS5 MTase in complex with HC. In addition, a fluorescence polarization-based assay was used to analyze the inhibitory effects of HC on GTP binding to DENV3 NS5 MTase. Therefore, the findings of this study offer a detailed understanding of the inhibition mechanism of HC.

## Methodology

### Fluorescence polarization

The fluorescence polarization (FP) assay was employed to measure the displacement of FAM-labeled GTP (F-GTP) binding to the DENV-3 NS5 methyltransferase (MTase) by HC. For K_D_ determinations, reaction mixtures were prepared with FP binding buffer (10 mM KCl,50 mM Tris, pH 7.5, 2 mM MgCl_2_, and 2 mM DTT , increasing concentrations of F-GTP (3.9 nM to 125 nM) and 250 nM NS5 MTase. For IC_50_ determinations of HC, the reaction mixtures contained FP binding buffer, increasing concentrations of HC (0 to 100 µM), 10 nM F-GTP, and 250 nM NS5 MTase. All samples were incubated in the dark at 25°C for 1 h before measuring total FP using a Synergy HTX multimode plate reader (Agilent BioTek) with Gen5 software, at an excitation wavelength of 485/20 nm and an emission wavelength of 528/20 nm. Baseline fluorescence polarization values from control samples containing only F-GTP (without protein) were subtracted to remove background signals. Data from three independent experiments were collected and analysed. K_D_ values were calculated using nonlinear regression analysis with a sigmoidal curve (18). IC_50_ values for HC were determined through nonlinear regression analysis, employing the log(inhibitor) versus normalized response curve (18), as analyzed by GraphPad Prism 8.0. The inhibition constant (Ki) for HC was calculated using the Cheng-Prusoff equation: K_i_= IC_50_/1+([F-GTP]/ K_D_) (18–20).

### Crystallization of NS5 MTase

DENV3 NS5 MTase was expressed and purified using recombinant plasmid pET28c(+)-NS5 MTase in *E. coli* BL21 cells, as described in the previous study (17). DENV3 NS5-MTase was crystallized employing the sitting drop vapor diffusion method in 96-well plates at 20°C, following the procedure outlined by Coutard et al., 2014 (17,21,22). To obtain the complex of DENV3 NS5-MTase with HC, soaking was done in cryoprotectant solution (22.5% glycerol) containing ∼ 1 mM HC for 30 min. The crystals were then flash-frozen in liquid nitrogen at 100K. The data collection of DENV3 NS5-MTase crystals in complex with HC was performed using Rigaku Micromax 007HF diffractometer integrated with Hypix 600C detector at the Macromolecular crystallography unit, IIT Roorkee. Data reduction and scaling were performed using the CrysAlis Pro PX software (Rigaku Inc.). The structure was solved by molecular replacement method using the MOLREP program of CCP4i2, giving DENV3 NS5 MTase (PDB code 4R8R) as a search model (23). Iterative rounds of model-building in COOT and the refinements were performed using the REFMAC5 (24) program in CCP4i2 (25). Non-crystallographic symmetry (NCS) and jelly-body restraints were applied during the entire data refinement process. Eventually, omit maps were generated in the Polder omit map program in Phenix software (26). Table 1 provides a summary of the data collection and refinement statistics. All figures of protein and ligand structures were prepared using PyMOL (27).

**Table 1.**
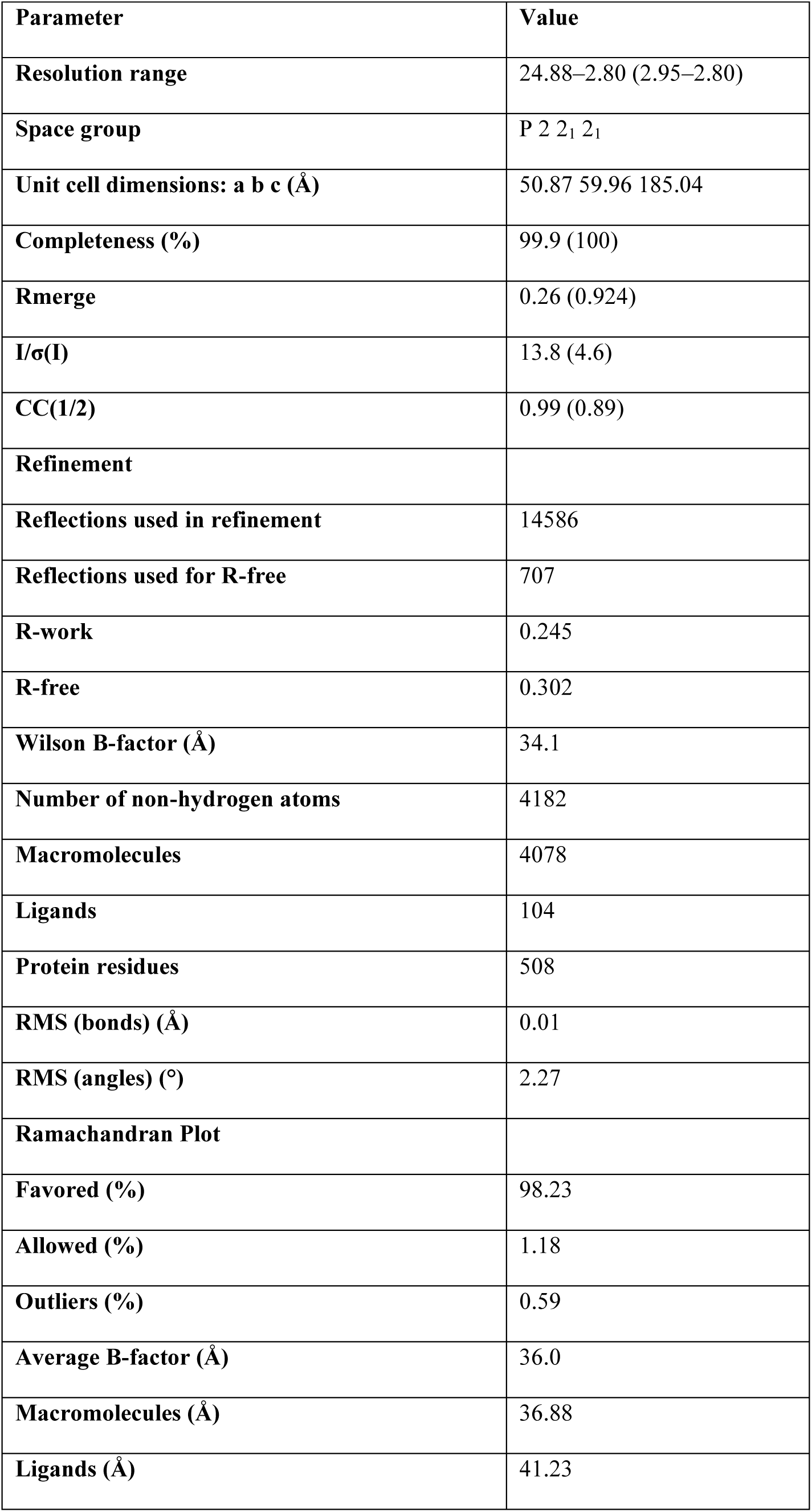
Data collection, processing, and refinement statistics of DENV3 NS5 MTase complexed with HC(PDB:8ZMC). Values in parentheses are for the highest-resolution shell. a Rmerge = Σ| I - 〈I〉|/ΣI. b R = Σ|Fobs| - |Fcalc|/Σ|Fobs|. The Rfree is the R calculated on the 5% reflections excluded for refinement. c RMS is root mean square.

## Results

### Inhibition of GTP binding to DENV3 NS5 MTase

A previous study demonstrated that HC binds to the recombinantly purified DENV3 NS5 MTase in the micromolar range K_D_ value. This binding was subsequently shown to exhibit anti-NS5 MTase activity. *In silico* work predicted that HC may bind within the GTP binding site (17). To investigate whether HC can displace GTP binding to DENV3 NS5 MTase, a fluorescence polarization (FP) assay was performed. To determine the dissociation constant (K_D_) of F-GTP, increasing concentrations of F-GTP were incubated with NS5 MTase. The binding curve revealed a sigmoidal relationship between F-GTP concentration and fluorescence polarization, resulting in a calculated K_D_ of ∼13.12 ± 1.02 nM (Figure 1A). This value indicates a high binding affinity of F-GTP for DENV3 NS5 MTase. A competition assay was conducted further to explore the inhibitory effect of HC on F-GTP binding. The inhibition curve demonstrated that HC effectively displaced F-GTP from the NS5 MTase, with a half-maximal inhibitory concentration (IC_50_) of 0.77 ± 0.01 µM (Figure 1B). Using the Cheng-Prusoff equation, the inhibition constant (K_i_) of HC was calculated to be approximately 0.43 ± 0.01 µM, suggesting that HC may inhibit GTP binding to DENV3 NS5 MTase.

**Figure 1:**
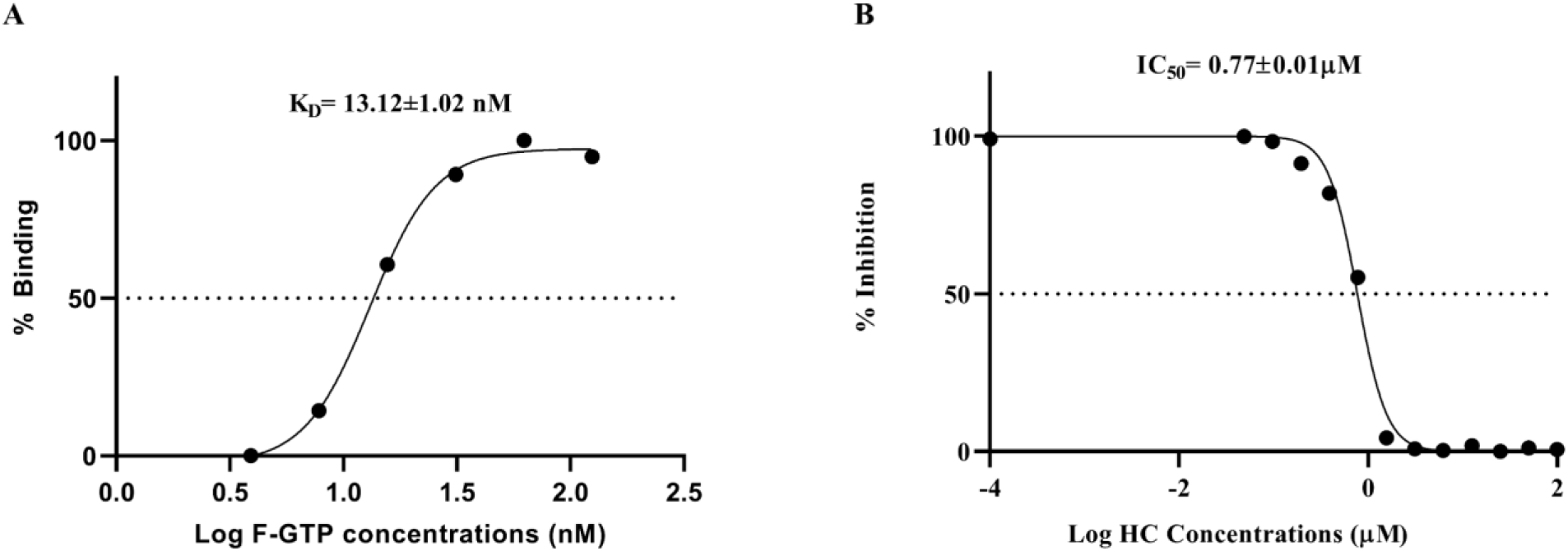
Fluorescence polarisation-based F-GTP displacement (A) Binding curve showing the interaction between F-GTP and DENV3 NS5 MTase. (B) Inhibition curve showing the effect ofHC on the binding of F-GTP to the DENV3 NS5 MTase.

### Structural insights into DENV3 NS5 MTase with HC

The structure of the DENV3 NS5 MTase in complex with HC (PDB: 8ZMC) was determined in the P 2 2_1_ 2_1_ space group at 2.8 Å with an R*_work_* of 0.24 and an R*_free_* of 0.30. The structure was solved using the molecular replacement method, with the apo form of DENV3 NS5 MTase (PDB ID: 4R8R) as a search model with 99.2 % identity. The refinement and data collection statistics are presented in Table 1. Structural analysis (Table 2) revealed the formation of hydrogen bonds and specific hydrophobic interactions, pivotal for the cap 0 RNA and GTP binding (27,28). One HC molecule was modeled in one chain of the DENV3 NS5 structure based on the difference Fourier map, which matched the observed electron density with a real space correlation coefficient (RSCC) of 0.78 (Figure 2B). The polder map contoured at 2.5 σ indicated its precise location within the DENV3 NS5 MTase (Figure 2A). The B-factor of HC was observed to be 34.1, which is similar to the B-factors of the nearby amino acids. The Ramachandran outliers were 0.59%, and none of the HC binding site residues contributed to this small fraction, thus proving the accuracy of the determined interaction of the binding site residues with HC (Table 1). In HC, O3 forms a hydrogen bond with the NZ of Lys42 at 2.85 Å, and O6 forms a hydrogen bond with the NE of Arg57 at a distance of 2.88 Å. HC hydrophobically interacts with Arg211, which is crucial for interacting with the phosphate group of GTP (Table 3) (Figure 2C). Although SAH was not added during crystallization, the electron density of SAH in the difference Fourier map was observed in both chains, suggesting that it would have been obtained from *E. coli* during protein expression (PDB:8ZMC) (Figure 2B). A similar observation of the electron density of SAH obtained from the host has been reported in previous studies (8,19) Additionally, SAH forms hydrogen bonds with the residues Asp131, Gly86, Ser56, Trp87, Lys105, and Val132. Hydrophobic interactions involve residues His110, Gly58, Gly81, Asp146, Lys130, Thr104, Gly83, Gly81, Asp146, Cys82, Gly85, and Phe133 (Figure 2D). The surface representation of the DENV3 NS5 MTase-HC superimposed with DENV3 NS5 MTase-SAH-cap 0RNA (PDB:5DTO) structures showed HC binds within the cap 0 RNA binding pockets (Figure 3A). The main chain conformation of the DENV3 NS5 MTase-HC complex closely resembles that of the DENV NS5-SAH-cap 0 RNA structure (PDB: 5DTO), with an RMSD of 0.354 Å across 233 Cα atoms (Figure 3B). In DENV3 NS5-SAH-cap 0 RNA, NH2 of Arg57 forms hydrogen bonds with OP1 of A-G (Table 4), while in the DENV3 NS5 MTase-HC complex, the O6 from HC forms hydrogen bonds with NE of Arg57, displaying binding of HC at the cap 0 RNA binding site (Table 3) (Figure 3D). The structural superposition of both complexes revealed that the HC interaction with Arg57 mediated angular variation of 124° the residue (Figure 3D). The superposition of DENV3 NS5 MTase bound to HC and GTP (PDB: 4V0R) demonstrates that HC forms hydrophobic interactions with Arg211, whereas GTP establishes a hydrogen bond with Arg211 (Figure 2C). These hydrophobic interactions of HC with the Arg211 residue of the GTP binding site conveyed a slight conformational change in Arg211. Additionally, Lys42 exhibits a notable angular shift of 51.6° in both complexes (Figure 3C).

**Figure 2:**
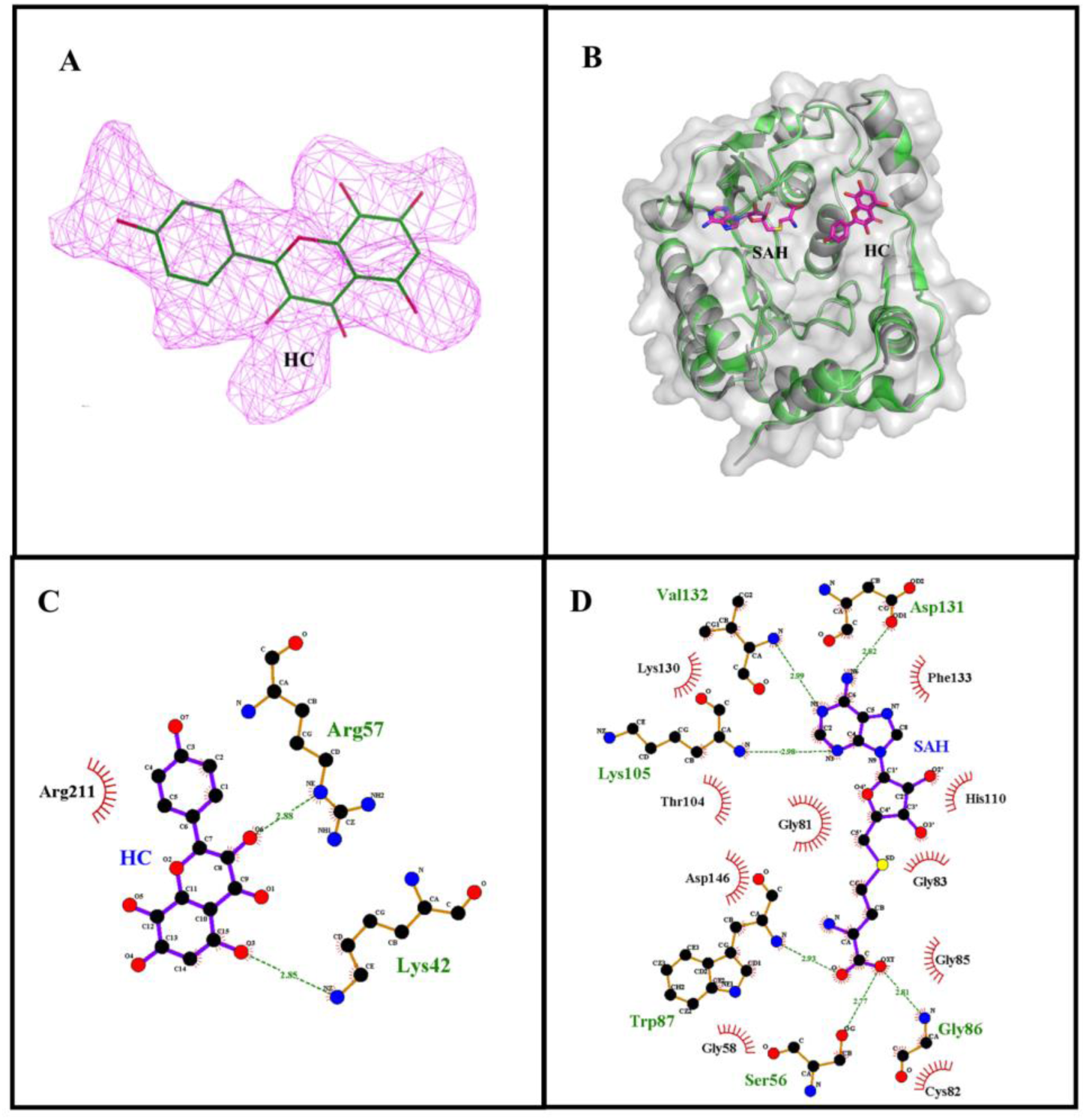
Structural analysis of the binding of HC and SAH to the DENV3 NS5 MTase. A) Polder omit map of HC at a contour level of 2.5σ. HC is displayed in green, and the polder map is shown as a magenta mesh. B) Surface and cartoon representation of the DENV3 NS5 MTase complex showing the binding pockets for HC and SAH. HC and SAH are depicted in stick representation, with HC and SAH in pink. The protein is shown in a green cartoon, and the surface is transparent grey. C) and D) 2D LigPlot representations for HC and SAH, respectively, showing the key residues involved in their interactions. Hydrogen bonds are depicted as green dashed lines, and hydrophobic contacts are represented by spiked red arcs.

**Figure 3:**
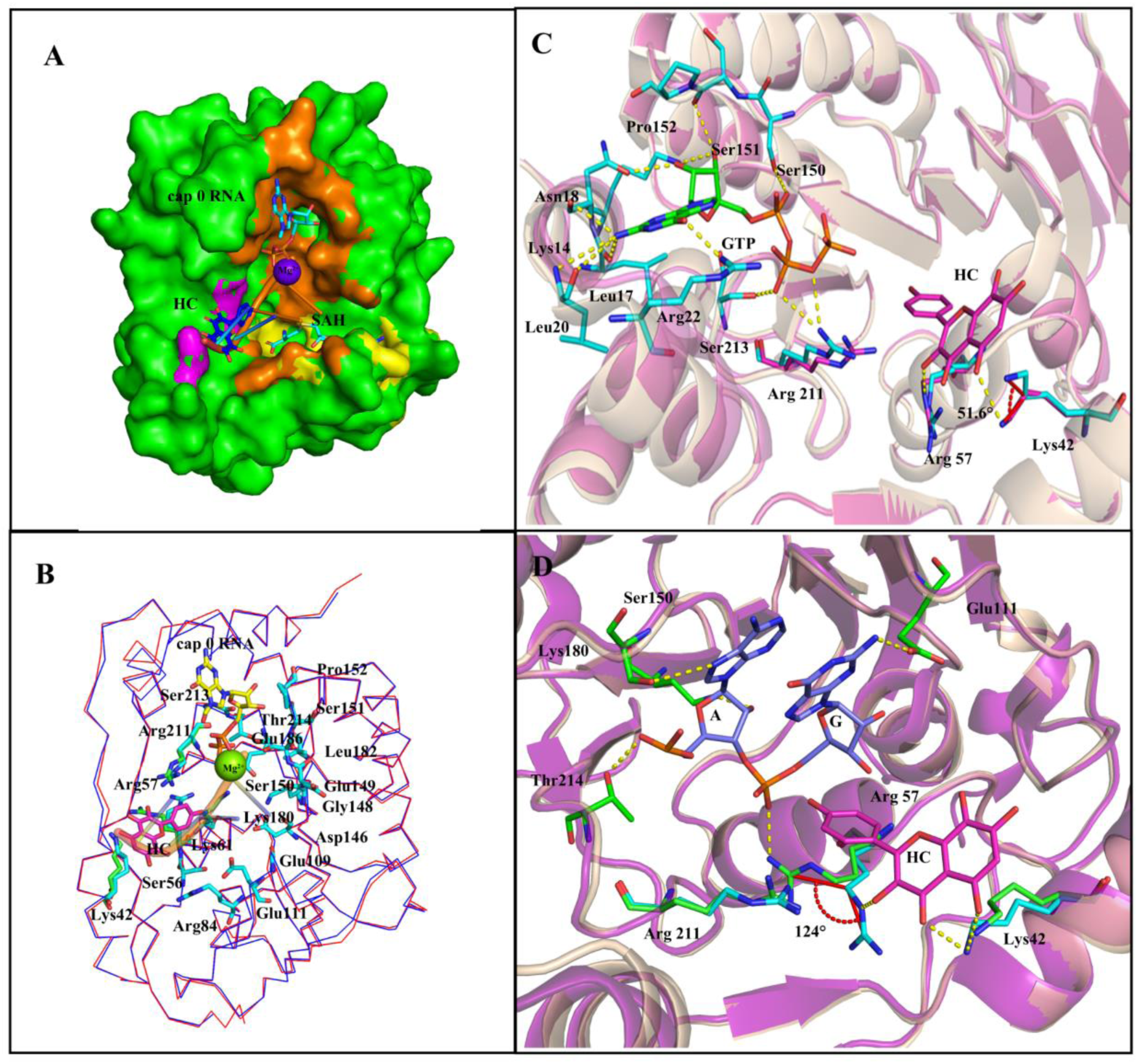
Structural analysis of HC, GTP, and cap 0 RNA interactions with DENV3 NS5 MTase. A) Surface representation of superposition of the DENV3 NS5 MTase-HC with DENV3 NS5 MTase-SAH-cap 0 RNA (PDB:5DTO). Binding pockets with Cap 0 RNA is shown in cyan, SAH in cyan, and HC in blue. Residues lining the cavity for cap 0 RNA are depicted in orange, SAH in yellow, and common residues from HC and cap 0 RNA in purple. A violet sphere indicates a bound Mg²⁺ ion. B) C-alpha chain superposition of DENV3 NS5 MTase-HC (blue) and the structure of GTP-bound MTase from PDB ID 5DTO (red). The ligands HC and cap 0 RNA are visualized with specific shapes and colors: cap 0 as a stick, RNA as a ribbon structure (yellow), and HC as a stick model (pink). All interacting residues are marked, with cap 0 RNA-interacting residues shown in blue and HC-interacting residues in green. The green sphere represents a bound Mg²⁺ ion. (C) Structure superposition of DENV3 NS5 MTase-HC (Pink-colored cartoon) with the structure from DENV3 NS5 MTase-GTP (PDB 4V0R) (wheat-colored cartoon). HC is represented in pink, with key residues interacting with HC also highlighted in pink. GTP is shown in green, residues are marked in blue (PDB 4V0R).D) Structure superposition of DENV3 NS5 MTase-A-G from cap 0 RNA (PDB 5DTO) (violet cartoon), with the DENV3 NS5 MTase-HC (wheat-colored cartoon). Adenine (A) and guanine (G) bases from the cap 0 RNA are colored blue, with residues highlighted in green (PDB 5DTO). HC is shown in pink, with residues highlighted in cyan. The red arc represents the angle of interaction. Hydrogen bonds between interacting residues are depicted as yellow dashed lines.

**Table 2:** Detailed molecular interactions between the DENV3 NS5 MTase and HC (PDB:8ZMC).

| Ligand | Interactions |  |  |  |
| --- | --- | --- | --- | --- |
|  | Hydrogen |  |  | Hydrophobic |
|  | Residues | Molecules | Bond length (Å) |  |
| HC | Arg57 | O6-NE | 2.88 | Arg211 |
|  | Lys42 | O3-NZ | 2.85 |  |

**Table 3:** Detailed molecular interactions between the DENV3 NS5 MTase domain and GTP (PDB:4V0R).

| Ligand | Interactions |  |  |  |
| --- | --- | --- | --- | --- |
|  | Hydrogen |  |  | Hydrophobic |
|  | Residues | Molecules | Bond length (Å) |  |
| GTP | Lys14 | O2'-NZ | 2.8 | Phe25,<br>Pro152,Asn18,<br>Lys29, Ser150 |
|  |  | O3'-NZ | 2.9 |  |
|  | Leu20 | N2-O | 3.4 |  |
|  | Asn18 | N2-O | 2.9 |  |
|  |  | O2'-OD1 | 3.0 |  |
|  | Ser151 | O3'-O | 3.3 |  |
|  | Arg22 | O6-NH1 | 2.6 |  |
|  | Leu17 | N2-O | 2.8 |  |
|  | Arg211 | O1B-NH1 | 3.3 |  |
|  |  | O3B-NH1 | 3.2 |  |
|  | Ser213 | O1B-OG | 3.1 |  |

**Table 4.**
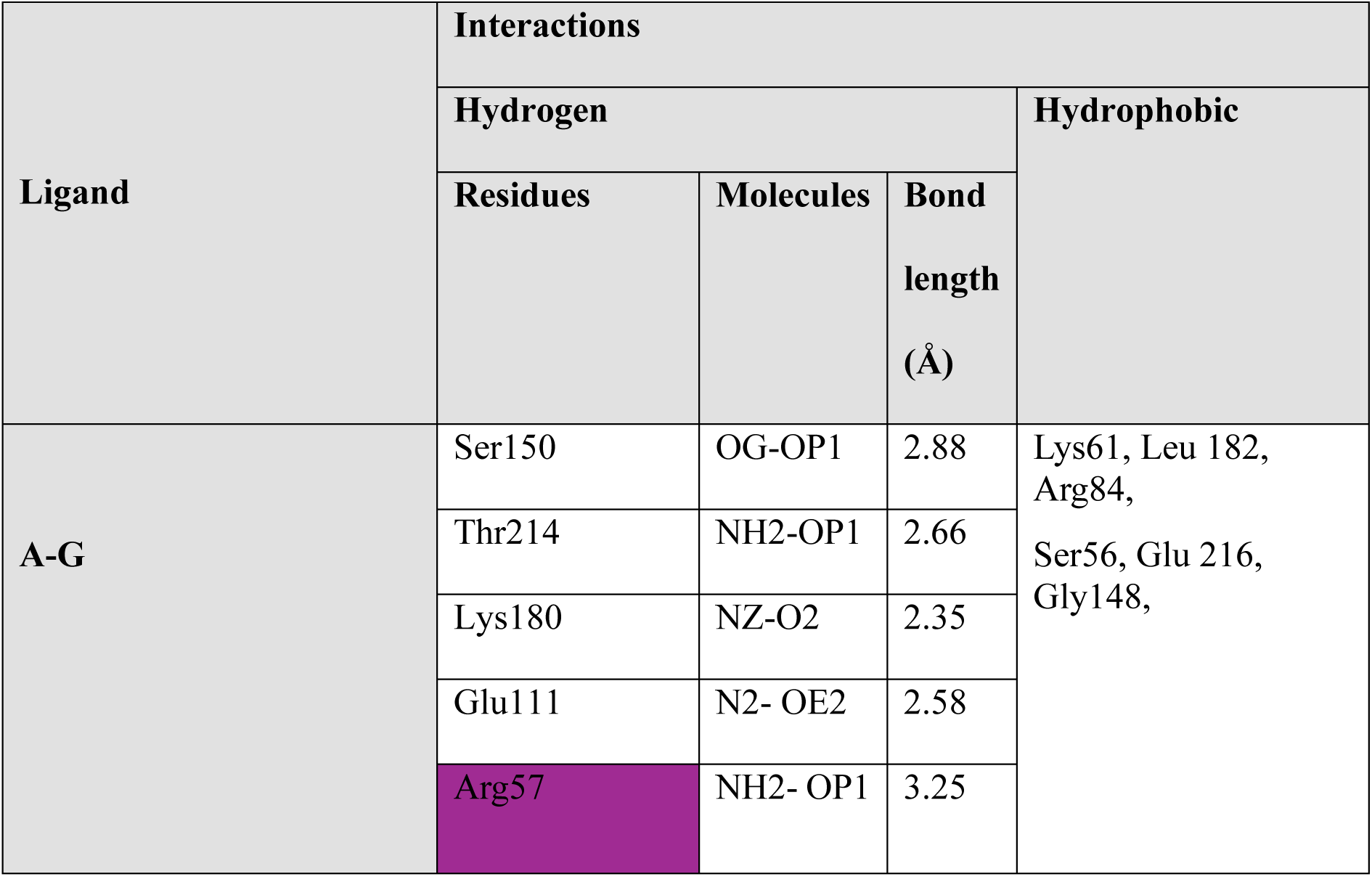
Detailed molecular interactions between the DENV3 NS5 MTase domain and A-G (from cap 0 RNA) (PDB:5DTO). These color-coded residues represent various interactions in a molecular context. Violet indicates a common residue that forms hydrogen bonds with HC and cap 0 RNA (A-G), and red signifies HC’s hydrophobic residues that make hydrogen bonds with GTP

| Ligand | Interactions |  |  |  |
| --- | --- | --- | --- | --- |
|  | Hydrogen |  |  | Hydrophobic |
|  | Residues | Molecules | Bond length (Å) |  |
| A-G | Ser150 | OG-OP1 | 2.88 | Lys61, Leu 182, Arg84, Ser56, Glu 216, Gly148, |
|  | Thr214 | NH2-OP1 | 2.66 |  |
|  | Lys180 | NZ-O2 | 2.35 |  |
|  | Glu111 | N2- OE2 | 2.58 |  |
|  | Arg57 | NH2- OP1 | 3.25 |  |

Consequently, the presence of HC at this site may obstruct the binding of GTP and viral RNA to the DENV3 NS5 MTase, resulting in effective structural inhibition of the enzyme.

### Conserved HC binding residues predicted to make stable interactions across DENV MTases of all serotypes

DENV 1-4, ZIKV, West Nile Virus (WNV), Yellow Fever Virus (YFV), and Japanese Encephalitis Virus (JEV) are orthoflaviviruses that cause infections in humans. Multiple sequence alignment (MSA) showed that the HC interacting residues Arg57, Lys42, and Arg211 remain conserved across all DENV serotypes (Supplementary Figure 1). Also, in other orthoflaviviruses such as ZIKV, JEV, and YFV, Lys42 is replaced by another positively charged amino acid Arg42, having similar chemical properties (Supplementary Figure 1). As the interacting residues (Arg57, Lys42, and Arg211) are conserved across all dengue serotypes, it is reasonable to hypothesize that HC would likely engage in similar key interactions with these residues across different serotypes. However, the non-interacting residues that vary between the serotypes could potentially influence the overall stability, orientation, or even the interaction dynamics of HC within the binding pocket. Leveraging this sequence conservation, HC coordinates from the DENV3 complex were superimposed onto the MTases of DENV1, 2, and 4. This superimposition was performed to determine the residues occupying structurally equivalent positions and interaction with the active site, as observed in the crystallized DENV3 MTase-HC complex, applied to all DENV serotypes (29).

Further, MD simulation studies were conducted to evaluate the interaction stability of MTase-HC complexes, with DENV3-HC MTase as the control for comparisons with other serotypes. The RMSD plots for DENV1, DENV2, DENV3, and DENV4 serotypes in complex with HC over the 100 ns MD simulation suggest stable binding interactions. The average RMSD values are 0.12 nm for DENV1, 0.13 nm for DENV2, 0.10 nm for DENV3, and 0.12 nm for DENV4 (Supplementary Figure 2A). The root mean square fluctuation (RMSF) data suggested that in MTase-HC complexes, the HC binding residues showed minimal fluctuations, indicating rigid and stable interactions with the ligand (Supplementary Figure 2 B). The radius of gyration (Rg) values for all four DENV serotypes exhibit a similar trend, indicating a stable conformation with average Rg values of 0.186 nm for DENV1, 0.187 nm for DENV2, 0.185 nm for DENV3, and 0.186 nm for DENV4 (Supplementary Figure 2C). The MTase-HC complexes maintained stable hydrogen bonds (H-bonds) throughout the simulation, with the average number of H-bonds being 1.30 for DENV1, 1.25 for DENV2, 1.77 for DENV3, and 0.73 for DENV4 (Supplementary Figure 2D).

The solvent accessible surface area (SASA) is key to understanding protein stability and folding DENV1, DENV3, and DENV4 complexes show comparable SASA values, whereas DENV2 exhibits slightly higher SASA values (Supplementary Figure 2E). The binding energies, van der Waals forces, and electrostatic interaction were calculated in terms of Lennard-Jones short-range (LJ-SR) and Coulombic SR (COU-SR), respectively. The LJ-SR values (in KJ/mol) are -71.6 for DENV1, -107.9 for DENV2, -75.15 for DENV3, and -32.09 for DENV4. The COU-SR values (in KJ/mol) are -60.75 for DENV1, -44.76 for DENV2, -51.88 for DENV3, and -27.24 for DENV4 (Supplementary Figure 2F). MD simulations of DENV serotypes reveal stable conformations over 100 ns, with subtle differences in structural dynamics and solvent exposure.

## Discussion

Previously, HC has been reported to exhibit antiviral efficacy against all DENV serotypes by targeting the viral replication enzyme NS5 MTase (17). This study has determined the crystal structure of DENV3 NS5 MTase in complex with the inhibitor HC to elucidate its inhibition mechanism. Additionally, the FP assay demonstrates that GTP is displaced due to the binding of HC (K_i_ value of ∼ 0.43 μM) to the conserved RNA binding site of NS5 MTase, which perturbs the spatial arrangement of GTP binding residues (Figure 3C).

The crystal structure of the HC-DENV3 MTase complex demonstrates that one HC molecule binds to the cap 0 RNA and near the GTP binding site (Figure 3). No differences were observed in the SAH molecule interactions compared to those reported in Zhao et al. 2015 (PDB: 4V0R) (8). GTP primarily establishes a hydrogen bonds with Lys14, Leu17, Asn18, Leu20, Arg22, Ser151, Arg211, and Ser213, residues and alongside forms multiple hydrophobic interactions involving Asn18, Phe25, Lys29, Ser150, and Pro152 residues (PDB:4V0R) (Table 3) (8). The RNA bases A-G form hydrogen bonds with Ser150, Thr214, Lys180, Glu111, and Arg57, and hydrophobic interactions with residues Leu182, Arg84, Lys61, Gly148, Ser56, and Glu216 (PDB:5DTO) (Table 4) (11). The 3-hydroxyphenyl ring of HC formed Pi-alkyl bond with Arg 57 while the trihydroxyphenyl ring formed Pi alkyl interaction with Lys42 revealing the binding of HC in the RNA binding site. Moreover, HC engages in hydrophobic interaction with Arg211. GTP forms two hydrogen bonds with Arg211, where one hydrogen bond is with the oxygen atom (O1B) of the second phosphate of GTP at a distance of 3.3 Å, and the other is with the oxygen atom (O3B), which links the second and third phosphates, at a distance of 3.2 Å (Table 3) (PDB: 4V0R). Thus, the interaction of HC with DENV3 MTase suggests its potential to display dual-mode inhibition of RNA and GTP binding.

Similarly, HC forms a hydrogen bond with Arg57 and Lys42, where Arg57 is a crucial residue that has been demonstrated to form hydrogen bonds with the viral RNA (PDB:5DTO) (11). The mutation Arg57Ala in the West Nile virus (WNV) MTase reduced N7 MTase activity to less than 20%(12). The detailed role of Lys42 is not well understood, but it may play an essential role in maintaining the structural stability of the NS5 MTase. Although this work did not investigate the inhibition of viral RNA binding to DENV3 NS5 MTase, HC’s interaction with key residues involved in RNA binding and reported MTase inhibition may hamper the viral RNA capping process (Table 2) (17). In MD simulations, the conserved HC binding residue is predicted to make stable interactions across the MTases of all DENV serotypes.

In conclusion, HC binds to the cap 0 RNA binding site, at the vicinity of the GTP binding site, thereby interfering with GTP binding. Further, *in vitro* and *in vivo* research is warranted to evaluate its efficacy across other orthoflaviviruses such as JEV, YFV, ZIKV, etc. Moreover, structural information and antiviral activity are anticipated to provide crucial insights for developing novel broad-spectrum inhibitors.

## Acknowledgments

The authors thank the Department of Biosciences and Bioengineering, IIT Roorkee for central facilities and the translational and structural bioinformatics center. The authors thank the Macromolecular Crystallographic Facility (MCU) at IIC, Indian Institute of Technology, Roorkee. ST and PK thank the Department of Biotechnology, Govt of India, for supporting Bioinformatics Center at IIT Roorkee ref number BT/PR40141/BTIS/137/16/2021. The authors thank the Ashok Soota Molecular Medicine facility, IIT Roorkee. ST and PK also would like to thank Department of Biotechnology, Govt of India “National Network Project of Department of Biotechnology, Indian Institute of Technology, Roorkee” Project no. BT/PR40142/BTIS/137/72/2023. The authors would like to thank the Council of Scientific and Industrial Research (CSIR), Indian Council of Medical Research (ICMR), and Ministry of Human Resource Development (MHRD) for providing financial support. This work was supported by a research grant to ST from DST SERB ref no. CRG/2023/002214

## Ethical approval

Ethical approval has been received for the work under the reference number IITR/IBSC/02/13/2024.

## Supplemental information

Supplemental information includes two supplemental experimental procedures and two Figures.

## Authors contribution

MB, SV, and VS conducted the experiments. MB, SV, VS, PK, and ST planned the experiments and analyzed the data. MB, PK, and ST took the lead in writing the manuscript. All authors provided critical feedback and helped shape the research, analysis, and manuscript.

## Supplementary Experimental Procedures

### Multiple Sequence Alignment (MSA)

The amino acid sequence of the NS5 MTase domain of flaviviruses was compared with the DENV3 NS5 MTase protein. DENV3 NS5 MTase were utilized as reference points for these alignments. The sequence alignment profile of NS5 MTase sequences was performed via Clustal Omega tool and analyzed by a graphical colored depiction using ESPript 3.0 (30). MSA was performed for the MTase domain of NS5 from DENV 1 (PDB:5IKM) , DENV 2 (PDB:1R6A), DENV3 (PDB: 5EHI), DENV 4 (Gene ID:NP_740325.1), Zika virus (ZIKV) (PDB:5WZ2), West Nile Virus (WNV) (PDB:2OY0), Yellow Fever Virus (YFV) (PDB:3EVA), and Japanese Encephalitis Virus (JEV) (PDB:4K6M) from the orthoflaviviruses and the analysis was carried out.

### MD simulation studies

The HC-bound MTase models were built using the experimentally resolved 3D structure of MTase (PDB: 8DKZ). The process involved aligning the MTase structures (PDB: 5IKM, 1R6A, and modelled structure of DENV4) to the template structure (PDB: 8DKZ), followed by the removal of other ligands and molecules after alignment (31). All procedures for constructing these HC-bound MTase models were performed using PyMOL. Subsequently, the prepared complexes underwent a 100 ns MD simulation to assess their stability and interactions. All simulation studies were performed using GROMACS 2022.2 (32) with the CHARMM36 force field (33) on a LINUX-based workstation. Ligand parameters and topology files were generated using the CHARMM General Force Field (CGenFF) program (34,35). The system was modelled in a cubic box, neutralized by adding ions and solvated with water molecules. Energy minimization was conducted using the steepest descent algorithm, followed by a two-phase equilibration for 100 ps, first under constant number of particles, volume, and temperature (NVT), and then under constant number of particles, pressure, and temperature (NPT). During NVT equilibration, the temperature was set to 300 K using a short-range electrostatic cut-off of 1.2 nm with the Berendsen thermostat (36). Afterward, the system transitioned to NPT equilibration, where coordinates were generated every 1 ps. A 100 ns production run was conducted with a 2fs time step, saving trajectories every 10 ps. The resulting trajectories were used to assess the stability and interactions of protein-ligand complexes.

---

**Supplemetnary Figure 1:**
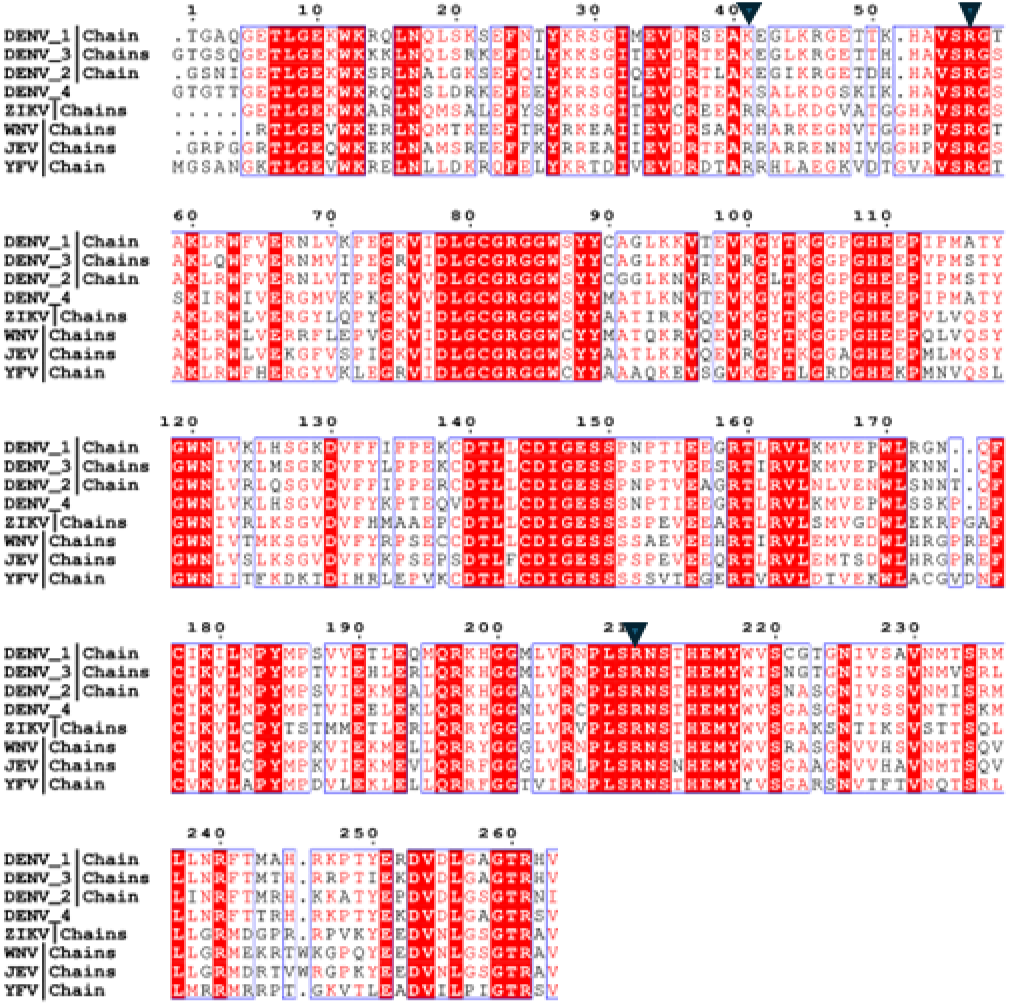
Sequence alignment of NS5 MTase domain (residues 1 to 263) from DENV with MTase domains of other flaviviruses. DENV 1 (PDB: 5IKM), DENV 2 (PDB:1R6A), DENV3 (PDB: 5EHI), DENV 4 (Gene id:NP_740325.1), ZIKV (PDB:5WZ2), WNV (PDB:2OY0), YFV (PDB:3EVA), and JEV (PDB:4K6M). Residues are represented with single letter amino acid code, with identical residues indicated in white font and boxed in red, while similar residues are indicated in red font. Blue triangles correspond to the HC binding site residues. Alignments were generated via Clustal Omega and ESPript tools.

**Supplementary Figure 2:**
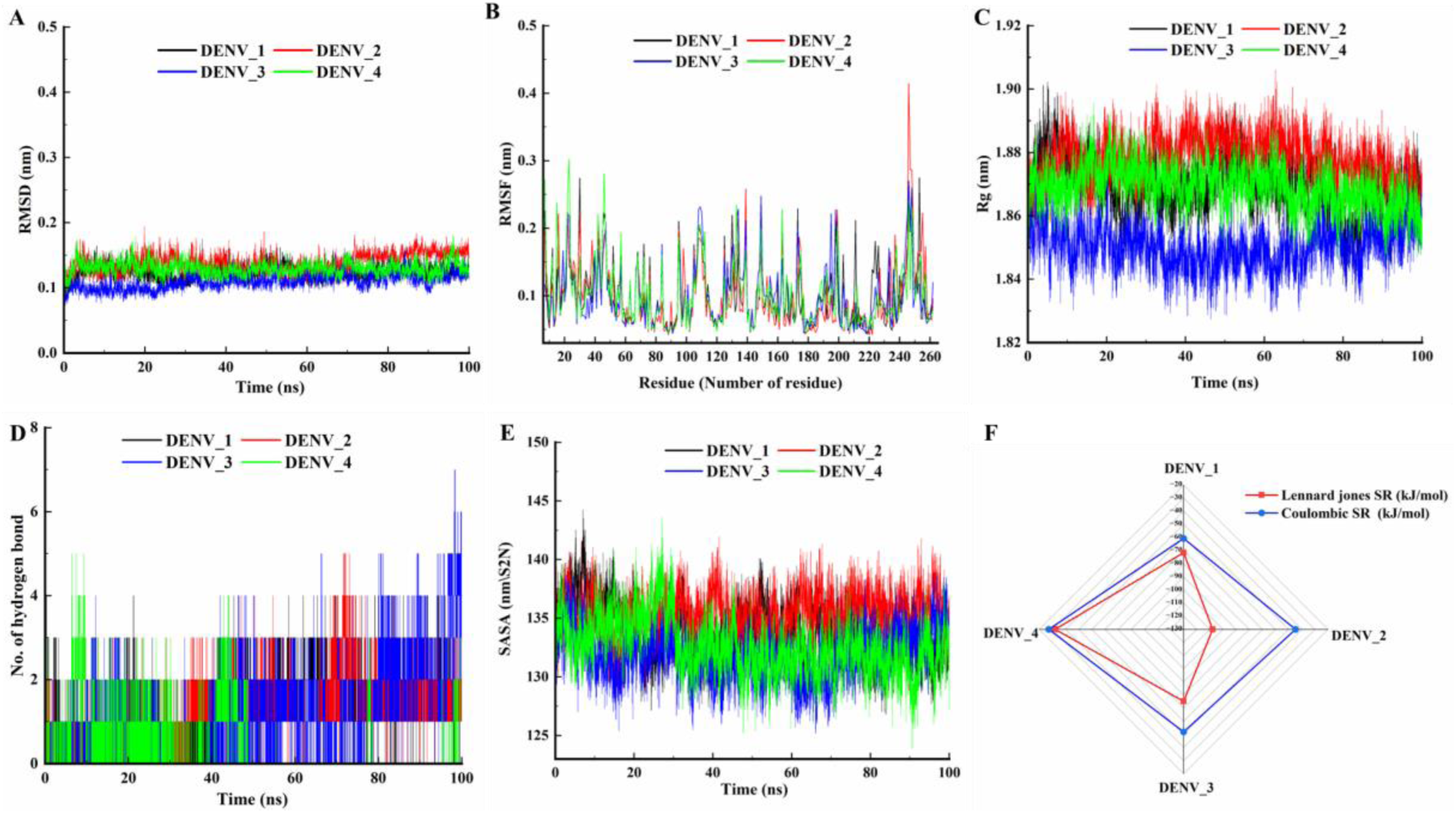
The MD analysis of MTase-HC complexes for DENV1, DENV2, DENV3, and DENV4 serotypes over a 100 ns simulation. A) RMSD plots of all DENV serotypes. B) RMSF data for all DENV serotypes with the black dashed circle indicating the HC binding residue. C) Rg values indicate conformations for all serotypes D) The hydrogen bond analysis of all DENV serotypes. E) SASA values of all DENV serotypes F). Radar plot showing the short-range (SR) interaction energies in kJ/mol. The red line represents the Lennard-Jones SR interaction energies, while the blue line represents the Coulombic SR interaction energies. Each vertex corresponds to a different serotype: DENV1, DENV2, DENV3, and DENV4.

## References

1. Postler TS, Beer M, Blitvich BJ, Bukh J, de Lamballerie X, Drexler JF, et al. Renaming of the genus Flavivirus to Orthoflavivirus and extension of binomial species names within the family Flaviviridae. Arch Virol. 2023;168(9):1–7. DOI: 10.1007/s00705-023-05835-1

2. Dengue and severe dengue. https://www.who.int/news-room/fact-sheets/detail/dengue-and-severe-dengue

3. Bhatt S, Gething PW, Brady OJ, Messina JP, Farlow AW, Moyes CL, et al. The global distribution and burden of dengue. 2013;496(7446):504–7. DOI: 10.1038/nature12060

4. Dengue-Global situation. https://www.who.int/emergencies/disease-outbreak-news/item/2023-DON498

5. Thomas SJ, Yoon IK. A review of Dengvaxia®: development to deployment. Hum Vaccin Immunother. 2019 (10):2295–314. DOI: 10.1080/21645515.2019.1658503

6. Murugesan A, Manoharan M. Dengue Virus. Emerging and Reemerging Viral Pathogens.2020 Jan 1;281. DOI: 10.1016/B978-0-12-819400-3.00016-8

7. Lim SP, Noble CG, Shi PY. The dengue virus NS5 protein as a target for drug discovery. Antiviral Res. 2015 ;119:57–67. DOI: 10.1016/j.antiviral.2015.04.010

8. Zhao Y, Soh TS, Zheng J, Chan KWK, Phoo WW, Lee CC, et al. A Crystal Structure of the Dengue Virus NS5 Protein Reveals a Novel Inter-domain Interface Essential for Protein Flexibility and Virus Replication. PLoS Pathog. 2015;11(3):1–27. DOI: 10.1371/journal.ppat.1004682

9. Furuichi Y, Shatkin AJ. Viral and cellular mRNA capping: Past and prospects. Adv Virus Res. 2000;55:135. DOI: 10.1016/s0065-3527(00)55003-9

10. Bisaillon M, Lemay G. Viral and Cellular Enzymes Involved in Synthesis of mRNA Cap Structure. Virology. 1997 ;236(1):1–7. DOI: 10.1006/viro.1997.8698

11. Zhao Y, Soh TS, Lim SP, Chung KY, Swaminathan K, Vasudevan SG, et al. Molecular basis for specific viral RNA recognition and 2’-O-ribose methylation by the dengue virus nonstructural protein 5(NS5). Proc Natl Acad Sci USA. 2015;112(48):14834–9. DOI: 10.1073/pnas.1514978112

12. Dong H, Ren S, Zhang B, Zhou Y, Puig-Basagoiti F, Li H, et al. West Nile virus methyltransferase catalyzes two methylations of the viral RNA cap through a substrate-repositioning mechanism. J Virol . 2008;82(9):4295–307. DOI: 10.1128/JVI.02202-07

13. Struijs K, Vincken JP, Doeswijk TG, Voragen AGJ, Gruppen H. The chain length of lignan macromolecule from flaxseed hulls is determined by the incorporation of coumaric acid glucosides and ferulic acid glucosides. Phytochemistry. 2009;70(2):262–9. DOI: 10.1016/j.phytochem.2008.12.015

14. Badshah SL, Faisal S, Muhammad A, Poulson BG, Emwas AH, Jaremko M. Antiviral activities of flavonoids. Biomedicine & Pharmacotherapy. 2021 ;140:111596. DOI: 10.1016/j.biopha.2021.111596

15. Rani R, Bhutkar M, Tomar S. Therapeutic Antiviral Potential of Flavonoids. In Flavonoids as Nutraceuticals 2024 Apr 9 (pp. 57–101). Apple Academic Press.

16. Kim DJ, Roh E, Lee MH, Oi N, Lim DY, Kim MO, et al. Herbacetin Is a Novel Allosteric Inhibitor of Ornithine Decarboxylase with Antitumor Activity. DOI: 10.1158/0008-5472.CAN-15-0442

17. Bhutkar M, Ruchi R, Kothiala A, Mahajan S, Waghmode B, Kumar R, et al. Elucidation of the antiviral mechanism of natural therapeutic molecules Herbacetin and Caffeic acid phenethyl ester against Chikungunya and Dengue virus. bioRxiv. 2022. DOI:https://www.biorxiv.org/content/10.1101/2022.05.31.494145v1

18. Bullard-Feibelman KM, Fuller BP, Geiss BJ. A Sensitive and Robust High-Throughput Screening Assay for Inhibitors of the Chikungunya Virus nsP1 Capping Enzyme. 2016. DOI: 10.1371/journal.pone.0158923

19. Lim SP, Sonntag LS, Noble C, Nilar SH, Ng RH, Zou G, et al. Small Molecule Inhibitors That Selectively Block Dengue Virus Methyltransferase. Journal of Biological Chemistry. 2011 Feb 25;286(8):6233–40. DOI: 10.1074/jbc.M110.179184

20. Dhaka P, Singh A, Choudhary S, Peddinti RK, Kumar P, Sharma GK, et al. Mechanistic and thermodynamic characterization of antiviral inhibitors targeting nucleocapsid N-terminal domain of SARS-CoV-2. Arch Biochem Biophys. 2023;750:109820. DOI: 10.1016/j.abb.2023.109820

21. Coutard B, Decroly E, Li C, Sharff A, Lescar J, Bricogne G, Barral K. Assessment of Dengue virus helicase and methyltransferase as targets for fragment-based drug discovery. Antiviral research. 2014 ;106:61–70. DOI: 10.1016/j.antiviral.2014.03.013

22. Benmansour F, Trist I, Coutard B, Decroly E, Querat G, Brancale A, et al. Discovery of novel dengue virus NS5 methyltransferase non-nucleoside inhibitors by fragment-based drug design. Eur J Med Chem. 2017;125:865–80. DOI: 10.1016/j.ejmech.2016.10.007

23. Vagin A, Teplyakov A. MOLREP: an Automated Program for Molecular Replacement. urn:issn:0021-8898. 1997;30(6):1022–5. DOI: 10.1107/S0907444909042589

24. Murshudov GN, Skubák P, Lebedev AA, Pannu NS, Steiner RA, Nicholls RA, et al. REFMAC5 for the refinement of macromolecular crystal structures. Acta Crystallogr D Biol Crystallogr. 2011 Apr;67(Pt 4):355–67. DOI: 10.1107/S0907444911001314.

25. Project CC. The CCP4 suite: programs for protein crystallography. Acta crystallographica. Section D, Biological crystallography. 1994 Sep 1;50(Pt 5):760–3. DOI: 10.1107/S0907444994003112

26. Liebschner D, Afonine PV, Moriarty NW, Poon BK, Sobolev OV, Terwilliger TC, Adams PD. Polder maps: improving OMIT maps by excluding bulk solvent. Acta Crystallographica Section D: Structural Biology. 2017;73(2):148–57. DOI: 10.1107/S2059798316018210

27. DeLano WL. The PyMOL Molecular Graphics System. DeLano Scientific. 2002

28. Laskowski RA, Swindells MB. LigPlot+: multiple ligand-protein interaction diagrams for drug discovery. J Chem Inf Model. 2011 ;51(10):2778–86. DOI: 10.1021/ci200227u.

29. Petrey D, Fischer M, Honig B. Structural relationships among proteins with different global topologies and their implications for function annotation strategies. Proceedings of the National Academy of Sciences. 2009;106(41):17377–82. DOI: 10.1073/pnas.0907971106

30. Sievers F, Wilm A, Dineen D, Gibson TJ, Karplus K, Li W, et al. Fast, scalable generation of high-quality protein multiple sequence alignments using Clustal Omega. Mol Syst Biol. 2011;7. DOI: 10.1038/msb.2011.75

31. Weng YL, Naik SR, Dingelstad N, Lugo MR, Kalyaanamoorthy S, Ganesan A. Molecular dynamics and in silico mutagenesis on the reversible inhibitor-bound SARS-CoV-2 main protease complexes reveal the role of lateral pocket in enhancing the ligand affinity. Scientific Reports 2021 11:1 . 2021;11(1):1–22. DOI: 10.1038/s41598-021-86471-0

32. Van Der Spoel D, Lindahl E, Hess B, Groenhof G, Mark AE, Berendsen HJC. GROMACS: fast, flexible, and free. J Comput Chem. 2005 ;26(16):1701–18. DOI: 10.1002/jcc.20291

33. Vanommeslaeghe K, Mackerell AD. CHARMM additive and polarizable force fields for biophysics and computer-aided drug design. Biochimica et Biophysica Acta (BBA) - General Subjects. 2015;1850(5):861–71. DOI: 10.1016/j.bbagen.2014.08.004

34. Vanommeslaeghe K, Hatcher E, Acharya C, Kundu S, Zhong S, Shim J, et al. CHARMM General Force Field: A Force Field for Drug-Like Molecules Compatible with the CHARMM All-Atom Additive Biological Force Fields. J Comput Chem. 2010;31:671–90. DOI: 10.1002/jcc.21367

35. Bhutkar M, Saha A, Tomar S. Viral methyltransferase inhibitors: berbamine, venetoclax, and ponatinib as efficacious antivirals against chikungunya virus. Arch Biochem Biophys. 2024;759:110111. DOI: 10.1016/j.abb.2024.110111

36. Berendsen HJC, Postma JPM, Van Gunsteren WF, Dinola A, Haak JR. Molecular dynamics with coupling to an external bath. J Chem Phys. 1984;81(8):3684–90. DOI:10.1063/1.448118

